# Independent parental contributions initiate zygote polarization in *Arabidopsis thaliana*

**DOI:** 10.1101/2020.12.02.407874

**Authors:** Kai Wang, Houming Chen, Yingjing Miao, Yanfei Ma, Agnes Henschen, Jan U. Lohmann, Sascha Laubinger, Martin Bayer

**Affiliations:** Max Planck Institute for Developmental Biology, Department of Cell Biology, Max-Planck-Ring 5, 72076 Tübingen, Germany; Department of Stem Cell Biology, Centre for Organismal Studies, Heidelberg University; 69120 Heidelberg, Germany; University of Oldenburg, Institute for Biology and Environmental Sciences, Carl-von-Ossietzky-Str. 9-11, 26111 Oldenburg, Germany

**Keywords:** *Arabidopsis thaliana*, MAP kinase signaling, ERECTA signaling, embryogenesis, BRASSINOSTEROID SIGNALING KINASE, SHORT SUSPENSOR, cell polarity, zygote, suspensor, parental conflict

## Abstract

Embryogenesis of flowering plants is initiated by polarization of the zygote, a prerequisite for correct axis formation in the embryo. Zygote polarity and the decision between embryonic and non-embryonic development in the daughter cells is controlled by a MITOGEN-ACTIVATING PROTEIN (MAP) kinase signaling pathway including the MAPKK kinase YODA (YDA). Upstream of YDA act two members of the BRASINOSTEROID SIGNALING KINASE (BSK) family, BSK1 and BSK2. These membrane-associated proteins serve as signaling relay linking receptor kinase complexes with intracellular signaling cascades. The receptor kinases acting upstream of BSK1 and BSK2 in the zygote, however, have so far not been identified. Instead, YDA can in part be activated by the non-canonical BSK family member SHORT SUSPENSOR (SSP) that is contributed by the sperm cell during fertilization. Here, we show that the receptor kinase ERECTA plays a crucial role in zygote polarization as maternally contributed part of the embryonic YDA pathway. SSP on the other hand provides an independent, paternal input to YDA activation, outlining a Y-shaped pathway. We conclude that two independent parental contributions initiate zygote polarization and control suspensor formation and embryonic development.

## Introductory paragraph

Embryogenesis of flowering plants is initiated by polarization of the zygote, a prerequisite for correct axis formation in the embryo. In Arabidopsis, the zygote elongates about three-fold before it divides asymmetrically. The smaller apical daughter cell forms the pro-embryo while the larger basal cell is the precursor of the mostly extra-embryonic suspensor ^1^. This filamentous organ plays a pivotal role in nutrient and hormone transport and is essential for the rapid growth of the embryo ^2^.

Zygote elongation and polarization is controlled by a MITOGEN-ACTIVATING PROTEIN (MAP) kinase signaling pathway including the MAPKK kinase YODA (YDA; ^3^). In *yda* mutants, the zygote divides symmetrically without obvious elongation. In many cases, the basal daughter cells follow an embryonic division pattern, yielding embryos with shorter or no suspensor. Constitutively active variants of YDA (*yda*-CA) give rise to a filament of daughter cells without recognizable pro-embryo ^3^. Therefore, this pathway seems to control embryonic vs non-embryonic development in the early embryo.

Upstream of YDA act two members of the BRASINOSTEROID SIGNALING KINASE (BSK) family, BSK1 and BSK2 ^4^. These membrane-associated proteins serve as signaling relay linking receptor kinase complexes with intracellular signaling cascades^5^. In other developmental processes, YDA activity is controlled by receptor kinases of the ERECTA family (ERf), including ERECTA (ER), ER-LIKE1 (ERL1), and ERL2 ^6–10^. In the embryonic YDA pathway, however, receptor kinases upstream of BSK1 and BSK2 have so far not been identified ^1^. Instead, YDA can in part be activated by the non-canonical BSK family member SHORT SUSPENSOR (SSP) by an unusual parent-of-origin effect ^4,11^. SSP protein accumulates transiently in the zygote after fertilization presumably translated from sperm cell-derived transcripts. The SSP protein represents a naturally occurring, constitutively active variant of BSK1 and activates YDA as soon as it is present in the zygote ^4^. SSP promotes suspensor formation and reinforces the stereotypic patterning of the early Arabidopsis embryo. Therefore, it has been speculated that SSP might be a paternal component of a parental tug-of-war controlling resource allocation towards the embryo ^2,11^.

Here, we show that the receptor kinase ERECTA plays a crucial role in zygote polarization as maternally contributed part of the embryonic YDA pathway. SSP on the other hand provides an independent, paternal input to YDA activation, outlining a Y-shaped pathway. We conclude that two independent parental contributions initiate zygote polarization and control suspensor formation and embryonic development.

## Main

The embryonic YDA pathway plays a central role in zygote polarization and the decision between embryonic versus non-embryonic development. It is however still unclear if this pathway is activated in response to extra-cellular signals. In many aspects of Arabidopsis development, YDA activity is controlled by receptor kinases of the ERECTA family^7,8,10^. As transcripts of *ER* are present in the zygote (Table S1; ^12^), a critical function of *ER* in controlling zygote polarity seems plausible. However, homozygous *er erl1 erl2* triple mutants are sterile and produce no embryos ^9^. To address a possible function of *ER* in early embryogenesis, we carefully examined homozygous single, double as well as segregating triple mutants of *ERf* genes. When comparing *er* mutants with wildtype, weak defects in zygote elongation, zygote polarization and suspensor length became apparent (Figure 1 and Figure S1). Furthermore, aberrant division plane orientation in the suspensor can be observed (Figure S2). These defects are hallmarks of reduced YDA activity ^3^ and mimic the phenotypes of *ssp* or *bsk1 bsk2* double mutants ^4,11^. The *erl1 erl2* double mutant did not show detectable differences to wildtype, indicating a minor role of *ERL1* and *ERL2* in early embryogenesis as long as a functional *ER* is present. In the absence of *ER*, however, further loss of *ERL1* or *ERL2* enhanced the mutant phenotype, suggesting that they can partially take over *ER* function (Figure 1 and Figure S1). This is reminiscent of *ERf* function during stomata development, where individual ERf members have by and large distinct signaling functions but can take over the role of others in their absence ^8,13–15^.

**Figure 1:**
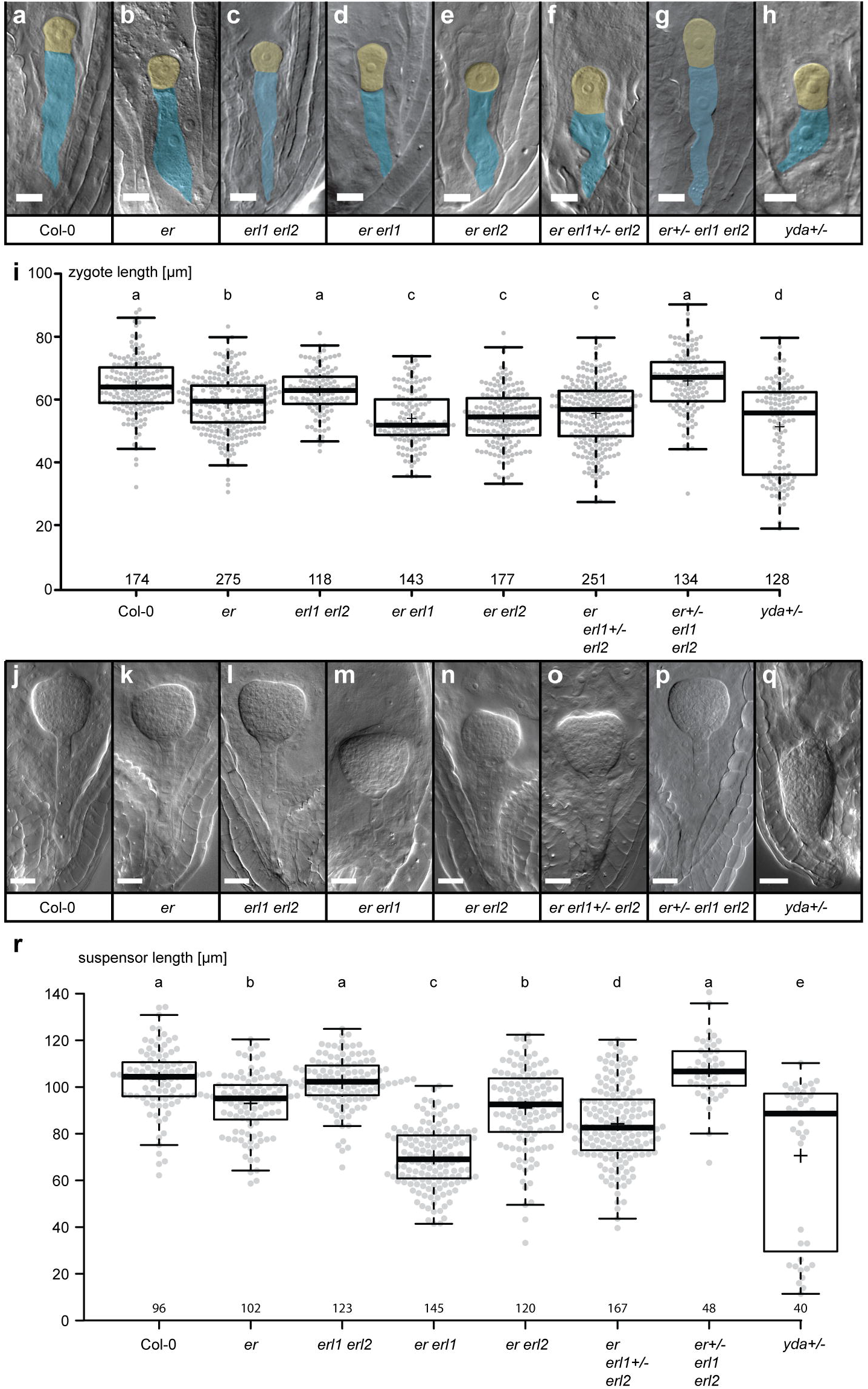
Zygote polarity and suspensor development. **a-h)** DIC images of cleared ovules showing representative 1-cell embryos. Apical cells are false colored in yellow, basal cells in blue. The genotype is given below the image. Heterozygous parental lines are indicated by +/−. Scale bar = 10μm. **i)** Box plot diagram of zygote length in μm. The sample size is given above x-axis. Center lines show the medians; box limits indicate the 25th and 75^th^ percentiles; whiskers extend 1.5 times the interquartile range from the 25th and 75th percentiles; crosses represent sample means; data points are plotted as grey dots. Statistical differences determined by Mann-Whitney U-test (p<0.05) are indicated by different letters above graph. In embryos of heterozygous *yda* plants, two distinct populations of data points can be observed, with approximately a quarter showing strong zygote elongation defects possibly representing the homozygous offspring. **j-q)** DIC images of cleared ovules showing transition stage embryos. The genotype is given below the image. Heterozygous parental lines are indicated by +/−. Scale bar = 20μm. **r)** Box plot diagram of suspensor length at transition stage. The sample size is given above the x-axis. Center lines show the medians; box limits indicate the 25th and 75th percentiles; whiskers extend 1.5 times the interquartile range from the 25th and 75th percentiles; crosses represent sample means; data points are plotted as grey dots. Statistical differences determined by Mann-Whitney U-test (p<0.05) are indicated by different letters above graph. In embryos of heterozygous *yda* plants, two distinct populations of data points can be observed, with approx. a quarter showing reduced suspensor length possibly representing the homozygous offspring.

To test if ER acts upstream of YDA in the context of zygote polarization as it does in other developmental contexts ^8,15,16^, we performed genetic rescue experiments with a constitutively active version of YDA (*yda-CA* ^3^; Figure S3). Constitutive YDA activity rescued the *er erl2* double mutant phenotype and resulted in a similar gain-of-function phenotype as observed in *yda-CA*, suggesting a function of *ER* and *YDA* in a common signaling cascade. Triple *ERf* mutants with segregating *er* (*er*+/− *erl1 erl2*), however, did not show defects in zygote elongation or suspensor formation despite normal transmission of the er mutant allele (Figure1, Table S2). The fact that phenotypic defects in homozygous embryos only became apparent if the parental plant was homozygous, indicated possible parent-of-origin effects. We tested this by analyzing F1 embryos of reciprocal crosses of *er erl* double mutants and wildtype (Figure 2 and Fig. S4). In these crosses, we could only observe embryonic defects if the maternal plant carried the *er erl* mutations but not if these alleles were transmitted by the pollen donor, indicating that ER function is under maternal control (Fig. 2, and Fig. S4). Furthermore, zygote elongation defects could not be observed if the maternal plant was heterozygous for the er mutation (Fig. 1), although slightly altered zygote polarity can be observed in the segregating triple *er +/− erl1 erl2* mutant (Fig. S1). The phenotype of the embryo therefore seems to be predominantly determined by the genotype of the maternal sporophyte while loss of *ER* function in the haploid phase or the embryo in segregating plants did not seem to have a major effect on the embryonic phenotype. This unusual parental effect may explain why a role of *ERECTA* in zygote polarization has so far been overlooked.

**Figure 2:**
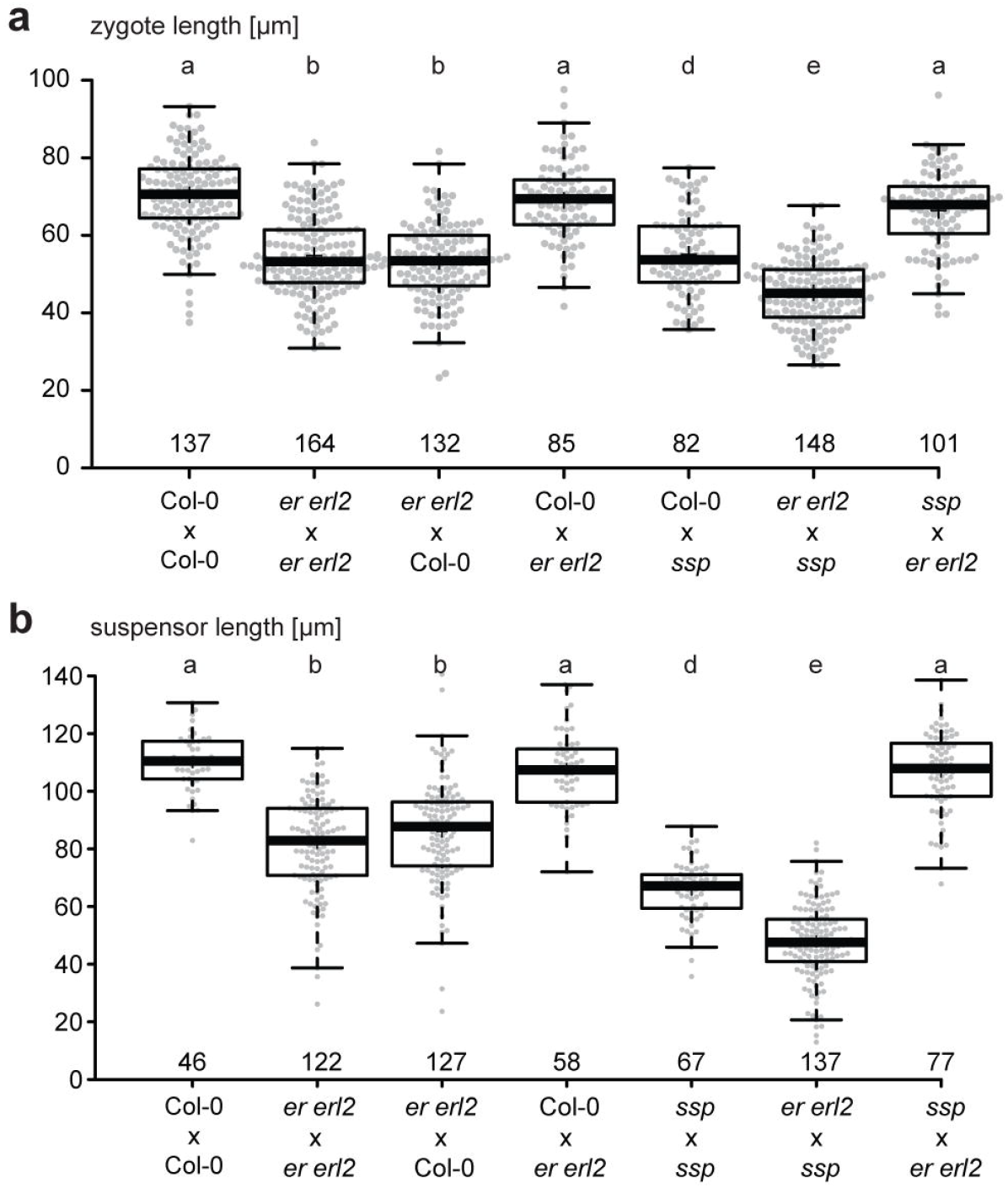
Parental effects in reciprocal crosses. Box plot diagrams of zygote elongation **a)** and suspensor length **b)** in μm in F1 embryos of reciprocal crosses. The embryonic defects of the *er erl2* double mutant depends solely on the maternal genotype. The paternal effect of *ssp* mutants enhances the *er erl2* defects in *er erl2* x *ssp*. The reciprocal *ssp* x *er erl2* cross does not show any detectable differences to wildtype. Genotypes are given as female x male. The sample size is indicated above the x-axis. Center lines show the medians; box limits indicate the 25th and 75th percentiles; whiskers extend 1.5 times the interquartile range from the 25th and 75th percentiles; small crosses represent sample means; data points are plotted as grey dots. Statistical differences determined by Mann-Whitney U-test (p<0.05) are indicated by different letters above graph.

Two possible scenarios could explain the sporophytic maternal effect: Zygote polarization could be affected by a non-cell autonomous function of ER in sporophytic tissue. Such a scenario has for example been described for early auxin signaling in the embryo ^17^. Alternatively, ER could function in the zygote but relying on *ER* transcripts and/or ER proteins inherited to the zygote via the pre-meiotic megaspore mother cell (MMC).

A transcriptional (*pER::3xVenus-N7*) and a functional translational fusion (*pER::ER-YPet*) reporter gene revealed *ER* expression in the sporophytic seed coat as well as the female germ line (Fig. S5). As *ER* transcripts can be detected in isolated egg cells and zygotes (Table S1; ^12^), we were not able to draw a definite conclusion about the location of ER function based on the observed expression pattern. However, *yda* is a zygotic recessive mutant ^3^ (Figure 1, Figure S1) and *SSP* acts as sperm-derived activator, arguing for a function of the embryonic ERECTA/YDA pathway in the zygote. When looking at mRNA stability, *ER* transcripts showed a moderate half-live in seedlings at roughly the same level as typical house-keeping genes, such as *ACTIN 2*, for which rapid mRNA turn-over does not appear necessary (Table S3)^18^. *ER* transcripts can be detected in the egg cell and early zygote but transcript levels decline after fertilization, indicating that there is little or no de novo transcription in the zygote^12^ (Table S1). In order to address the question where ER function is necessary for zygote polarization, we specifically reduced ER protein function in the zygote using the *pER::ER-YPet* line in *er erl1 erl2* background. Genetically encoded anti-GFP nanobodies fused to ubiquitin ligases have been used as elegant tool to specifically target GFP/YFP-tagged proteins for degradation ^19,20^. To first test its functionality, we expressed the NSlmb-vhhGFP4 nanobody under the *ER* promoter. The resulting seedlings mimicked the *er erl1 erl2* triple mutant phenotype albeit with a slightly weaker phenotype (Fig. 3), indicating that the nanobody is functional but does not completely abolish ER-YPet activity. In a next step, we expressed the nanobody under the strong egg cell-specific *EC1* promoter to specifically reduce ER-YPet levels in the egg cell/zygote. This resulted in a significant reduction of zygote length in the *pER::ER-YPet* rescue line, indicating that zygote polarization critically depends on functional *ER* protein in the zygote. Taken together, the available data suggests that the maternal control of ER function can be explained by the inheritance of pre-meiotically produced *ER* transcripts to the zygote, compensating the loss of *ERECTA* in the female germline.

**Figure 3:**
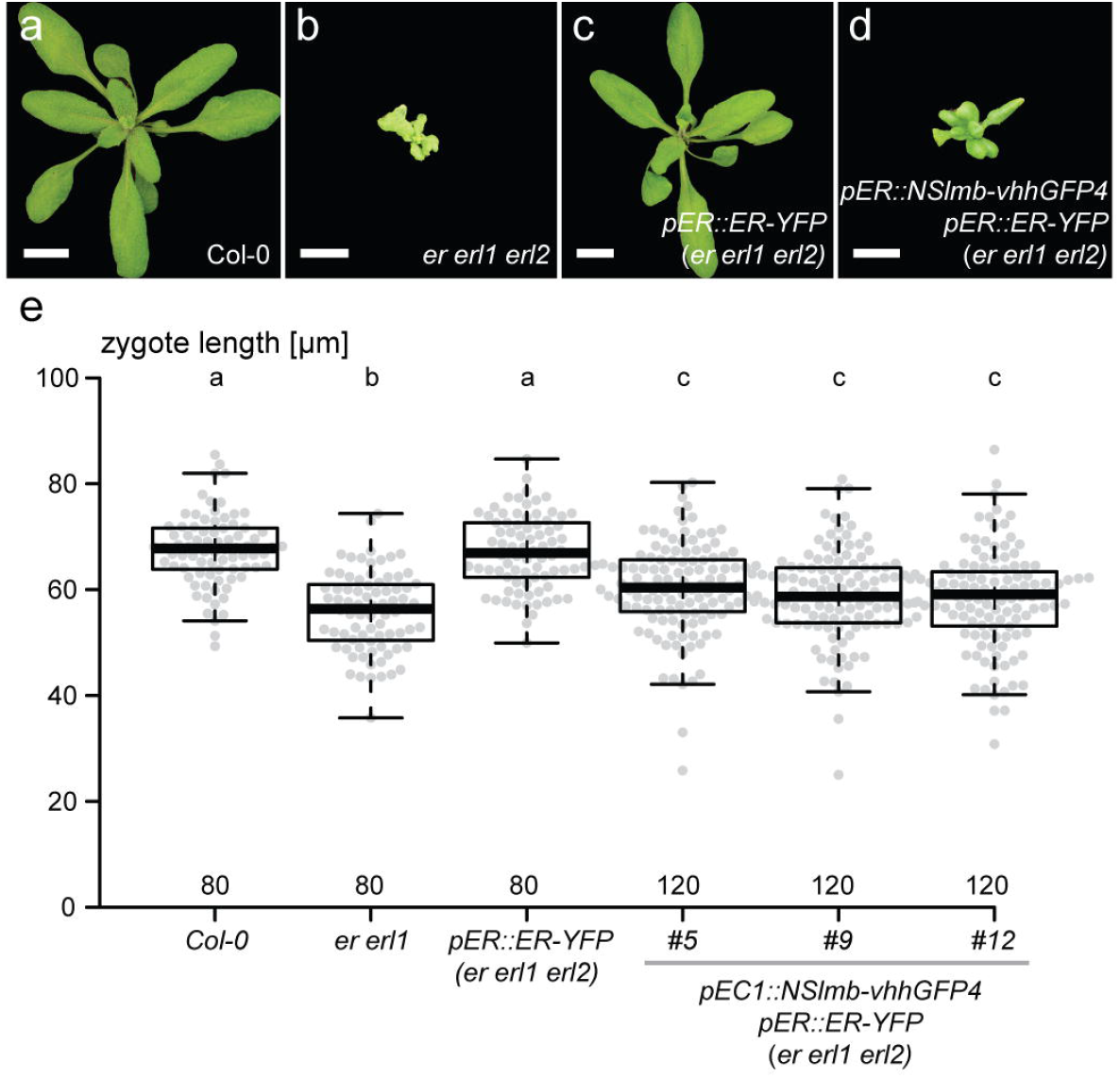
Cell-autonomous function of ER protein in the zygote. Vegetative growth phenotype of wild-type Col-0 **(a)**, *er erl1 erl2* triple homozygous **(b)**, genetically rescued plants (*pER::ER-YPet* in *er erl1 erl2*; **c)**, and genetically recued plants expressing anti-GFP nanobody (*pER::NSlmb-vhhGFP4* in *pER::ER-YPet er erl1 erl2*; **d)**. Scale bar = 1 cm. **e)** Egg cell-specific expression of the anti-GFP nanobody (*pEC1::NSlmb-vhhGFP4*) in the *pER::ER-YPet er erl1 erl2* plant leads to reduced zygote length. Box plot diagram of zygote elongation in μm. The sample size is indicated above the x-axis. Center lines show the medians; box limits indicate the 25th and 75th percentiles; whiskers extend 1.5 times the interquartile range from the 25th and 75th percentiles; small crosses represent sample means; data points are plotted as grey dots. Statistical differences determined by Mann-Whitney U-test (p<0.05) are indicated by different letters above graph.

BSK1 and BSK2 have been shown to act upstream of YDA, presumably linking ERECTA signaling with YDA activation ^4^. Reciprocal crosses with *bsk1 bsk2* double mutants and wildtype showed similar sporophytic maternal effects as observed for *er* (Figure S7). In addition, maternal effects have been described for mutations in genes acting downstream of YDA ^21,22^. Therefore, not just *ER* but several components of this signaling pathway appear to be under maternal control. This is quite intriguing, as the noncanonical BSK family member SSP activates YDA as paternal component contributed by the sperm cell ^11^. SSP has been shown to resemble a naturally occurring, constitutively active form of BSK that directly interacts with YDA via its TPR motif ^4^. This raises the question whether the two parental contributions to YDA activation can function independently. We tested this hypothesis in seedlings, where ectopic expression of *SSP* leads to a strong gain-of-function phenotype by constitutive activation of the YDA pathway ^4,11^. Ectopic expression of SSP-YFP (*p35S::SSP-YFP*) in wildtype resulted in seedlings with cotyledons lacking mature stomata (Figure 4). This was also observed for *SSP-YFP* expression in *er erl1 erl2* triple mutant background, indicating that SSP can activate YDA in the absence of a functional receptor complex.

**Figure 4:**
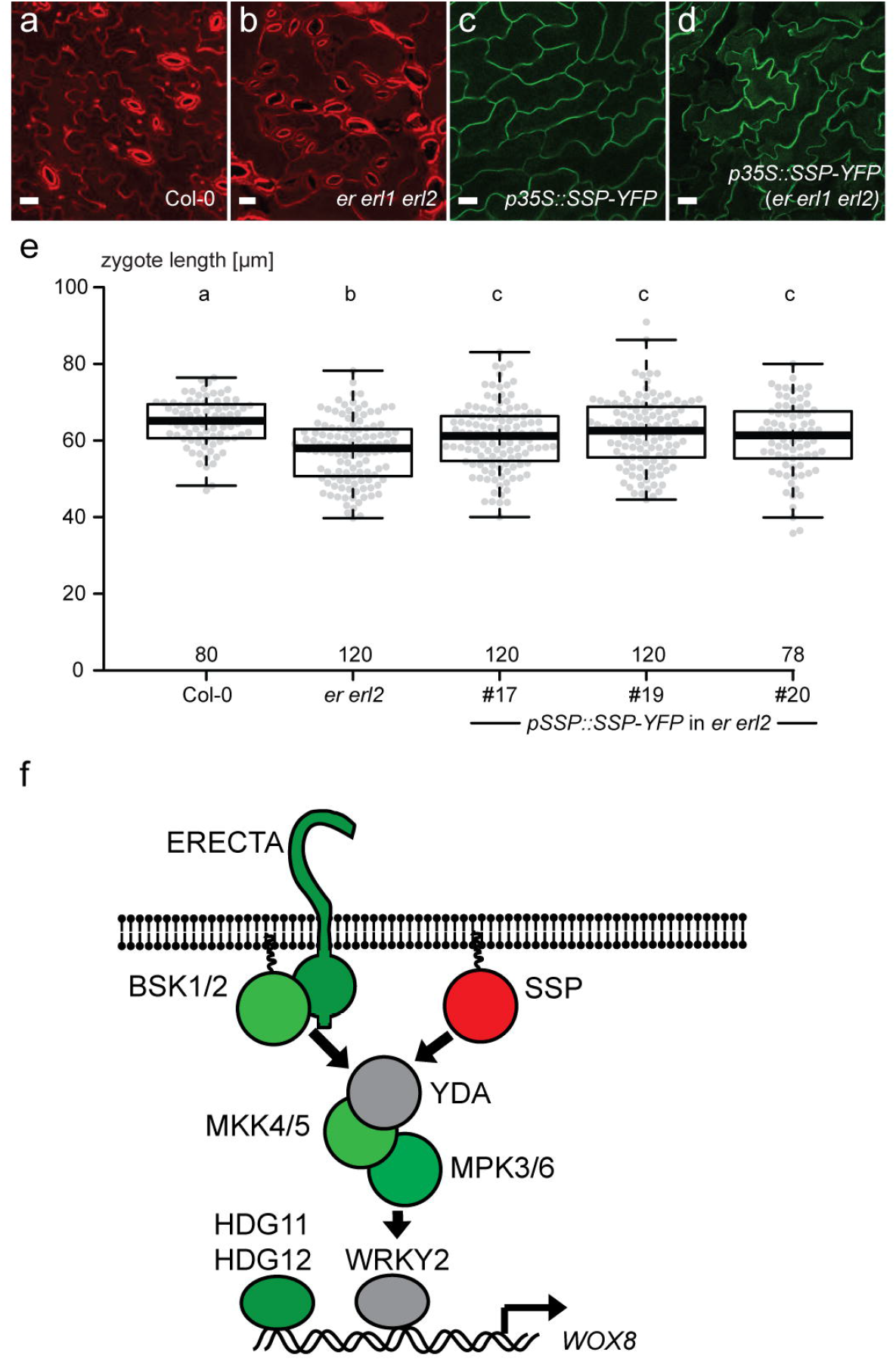
Y-shaped topology of the embryonic YDA pathway. Ectopic expression of *SSP-YFP* in leaves under control of the CaMV 35S promoter (*p35S::SSP-YFP*) leads to absence of stomata on the leaf surface in wild-type **(c)** and *er erl1 erl2* background **(d)**. Col-0 **(a)** and *er erl1 erl2* **(b)** are shown as reference. SSP-YFP signal is shown in green, propidium iodide staining in red. Scale bar = 20μm. An additional copy of SSP (*pSSP::SSP-YFP*) partially rescues the *er erl2* loss-of-function phenotype in the zygote **(e)**. Box plot diagram of zygote elongation in μm. The sample size is indicated above the x-axis. Center lines show the medians; box limits indicate the 25th and 75th percentiles; whiskers extend 1.5 times the interquartile range from the 25th and 75th percentiles; small crosses represent sample means; data points are plotted as grey dots. Statistical differences determined by Mann-Whitney U-test (p<0.05) are indicated by different letters above graph. **f)** Simplified schematic depiction of the embryonic YDA pathway. YDA is activated by the maternally supplied receptor complex consisting of ERECTA and the membrane-associated proteins BSK1 and BSK2. YDA can also be activated by the paternally contributed BSK family member SSP. Signaling components are colored according to genetic contribution: maternal effect (green), paternal effect (red), zygotic recessive (grey).

If SSP indeed serves as independent signal for YDA activation in the zygote, loss of *SSP* should have an additive effect to loss of *ER* function. We therefore observed the zygotic phenotype of *ssp, er erl*, and *er erl ssp* mutants (Figure 2 and Fig. S8). The further loss of *SSP* in *er erl1* mutant lead to further reduction in zygote length and an almost complete loss of zygote polarity. Taken together, this indicates that *SSP* provides additional YDA activation that is independent of *ER* function. If the embryonic YDA pathway indeed relies on two independent signaling inputs, additional SSP activity should compensate for the loss of ER function. To test this, we introduced an additional copy of SSP (pSSP::SSP-YFP) in the er erl2 double mutant and found that additional *SSP* activity indeed partially rescued the loss-of-function phenotype of *er erl2* (Figure 4).

Taken together, our data outlines a Y-shaped signaling pathway (Figure 4e) with a maternally controlled receptor complex and a paternally provided activating protein that converge at the level of MAP3K activation. *SSP* is a *Brassicaceae*-specific gene while *ERECTA, BSK1*, and *BSK2* are evolutionarily conserved in flowering plants ^23,24^. *SSP* has been shown to be critical for correct suspensor development, an organ that serves as conduit for nutrients to the embryo, and for rapid development of the embryo ^2^. Therefore, suspensor development could be a possible target of a parental conflict over nutrient allocation to the embryo. It has recently been observed that there are preferentially maternal expressed genes in the suspensor while there is mainly biparental expression in the apical cell ^25^. This maternal bias in the suspensor could be interpreted in the context of the parental conflict theory enforcing equal nutrient distribution to all embryos by the maternal genome^26^. In this scenario, *SSP* could be seen as a *Brassicaceae*-specific, paternal contribution that circumvents the maternal control over suspensor development by bypassing ERECTA signaling as strong activator of the YDA pathway.

This raises the question if suspensor development in other plant species similarly is also under maternal control and if other parental contributions exist. It will be fascinating to elucidate whether the two independent inputs for YDA activation indeed are part of a parental tug-of-war over nutrient allocation to the embryo or if *Brassicaceae* are shifting from ligand-/receptor-based zygote polarization to a sperm-based activation mechanism via SSP to fine-tune the timing of YDA activation as SSP naturally links fertilization with YDA activation.

## Acknowledgements

We would like to thank Gerd Jürgens for helpful discussions and comments on the manuscript. We also thank the Salk Institute Genomic Analysis Laboratory as well as the Nottingham Arabidopsis Stock Centre (NASC) for providing the sequence-indexed Arabidopsis T-DNA insertion mutants. Research in our groups is supported by the German Research Foundation (Deutsche Forschungsgemeinschaft - DFG SFB1101/B01 to M.B. and SFB1101/B07 to JUL) and the Max Planck Society.

## Author contributions

KW and MB designed the study with critical comments and modifications by HC, YMi, YMa, JUL, and SL. KW, HC, YMi, and AH performed the experiments. KW, HC, YMi, YMa, AH, JUL, SL, and MB evaluated the data and interpreted the experimental results. KW and MB wrote the initial manuscript draft, all authors commented on and edited the manuscript.

## Supplementary data

<u>Suppl. Figure S1:</u> Zygote polarity determined as ratio of apical and basal cell sizes

<u>Suppl. Figure S2:</u> Frequency of aberrant division planes in the suspensor

<u>Suppl. Figure S3:</u> Genetic rescue with *yda*-*CA*

<u>Suppl. Figure S4:</u> Sporophytic maternal effect of er erl1 double mutant

<u>Suppl. Figure S5:</u> *ERECTA* expression in developing ovules

<u>Suppl. Figure S6:</u> Expression of *pER::ER-YPet* in *er erl1 erl2*

<u>Suppl. Figure S7:</u> Sporophytic maternal effect of *bsk1 bsk2* double mutants

<u>Suppl. Figure S8:</u> Parental effects of *er erl1* and *ssp* mutations.

<u>Supplementary Table S1:</u> mRNA abundance of *ERECTA* family genes during early embryogenesis.

<u>Supplementary Table S2:</u> Segregation rates of *ER* and *ERL1* in triple mutant combinations

<u>Supplementary Table S3:</u> mRNA decay of selected candidate genes

<u>Supplementary Table S4:</u> primers and oligo nucleotide sequences

<u>Material and Methods</u>

