## Supplementary data and Methods for "Independent parental contributions initiate zygote polarization in *Arabidopsis thaliana*"

**Affiliations**

**Supplementary data:**

Suppl. Figure S1: Zygote polarity determined as ratio of apical and basal cell sizes

Suppl. Figure S2: Frequency of aberrant division planes in the suspensor

Suppl. Figure S3: Genetic rescue with *yda-CA*

Suppl. Figure S4: Sporophytic maternal effect of er erl1 double mutant

Suppl. Figure S5: *ERECTA* expression in developing ovules

Suppl. Figure S6: Expression of *pER::ER-YPet* in *er erl1 erl2*

Suppl. Figure S7: Sporophytic maternal effect of *bsk1 bsk2* double mutants

Suppl. Figure S8: Parental effects of *er erl1* and *ssp* mutations.

Supplementary Table S1: mRNA abundance of *ERECTA* family genes during early embryogenesis.

Supplementary Table S2: Segregation rates of *ER* and *ERL1* in triple mutant combinations

Supplementary Table S3: mRNA decay of selected candidate genes

Supplementary Table S4: primers and oligo nucleotide sequences

Material and Methods

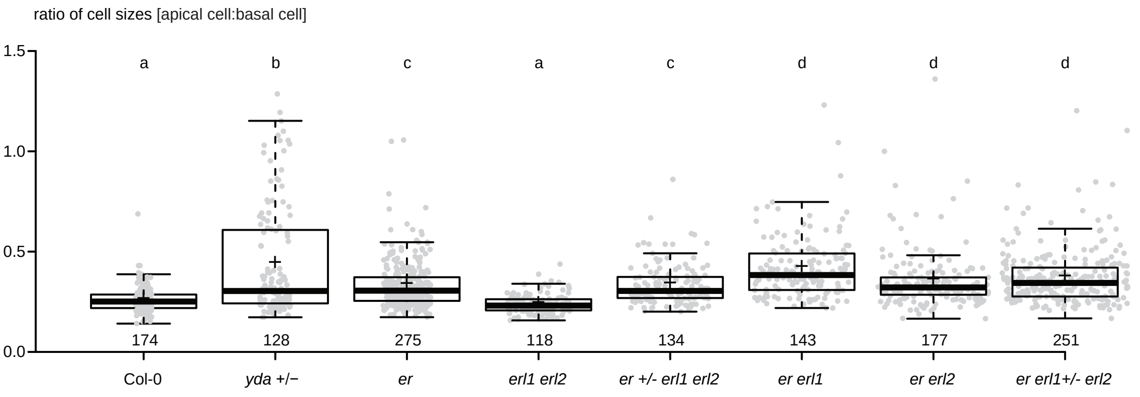

**Figure S1: Zygote polarity determined as ratio of apical and basal cell sizes**
Box plot diagram showing the ratio of apical to basal cell size (data of Fig. 1i). Center lines show the medians; box limits indicate the 25th and 75th percentiles; whiskers extend 1.5 times the interquartile range from the 25th and 75th percentiles; data points are plotted as grey dots. Statistical differences determined by Mann-Whitney U-test ($p<0.05$) are indicated by different letters above graph. In segregating offspring of heterozygous *yda* plants, two distinct populations of data points can be observed, with approx. a quarter showing strong zygote polarity defects possibly representing the homozygous offspring.

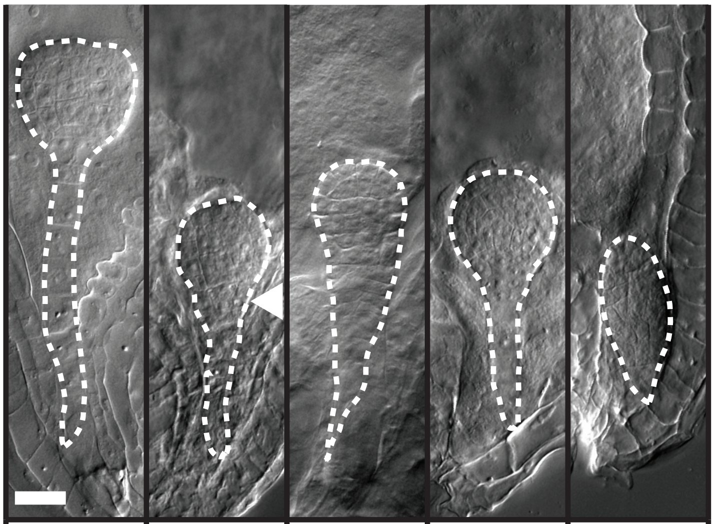

| Col-0 | *ssp* | *bsk1 bsk2* | *er erl1* | *yda+/-* |
|---|---|---|---|---|
| Frequency of aberrant division plane orientation | | | | |
| 0/242 | 14/90 | 12/88 | 5/107 | 35/155 |

**Figure S2: Frequency of aberrant division plane orientation in late globular stage embryos**
DIC images of cleared ovules showing late globular stage embryos. The genotype is given below the image. Aberrant division planes at the boundary of embryo proper and suspensor in the *ssp* sample are highlighted by an arrowhead. Heterozygous parental lines are indicated by +/-. Scale bar = 20 µm.

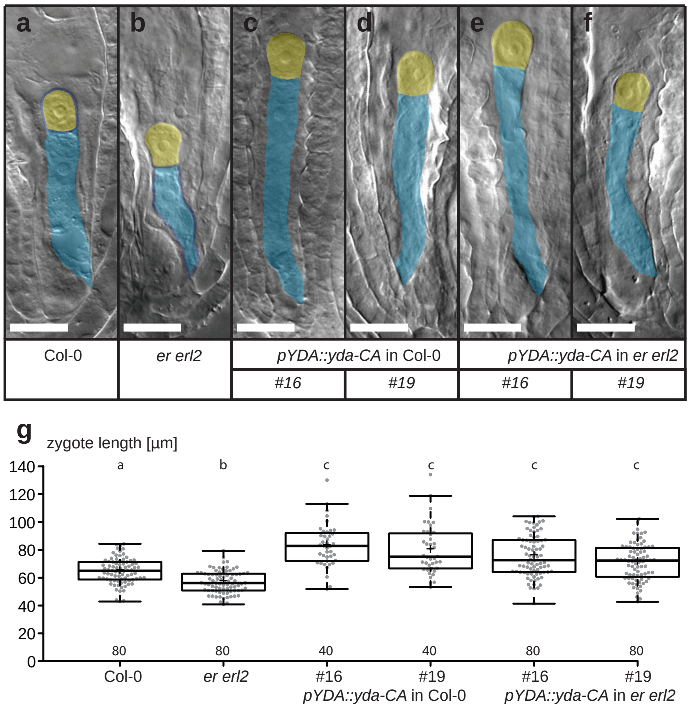

**Figure S3: Genetic rescue with *yda-CA***
DIC images of cleared ovules showing representative 1-cell embryos **(a-f)**. Apical cells are false colored in yellow, basal cells in blue. The genotype is given below the image. Two independent transgenic lines of *pYDA::yda-CA* in *er erl2* were used. As control, the same lines were backcrossed to Col-0. Scale bar = 20µm.
**g)** Box plot diagram of zygote length in µm. The sample size is given above x-axis. Center lines show the medians; box limits indicate the 25th and 75th percentiles; whiskers extend 1.5 times the interquartile range from the 25th and 75th percentiles; crosses represent sample means; data points are plotted as grey dots. Statistical differences determined by Mann-Whitney U-test ($p<0.05$) are indicated by different letters above graph.

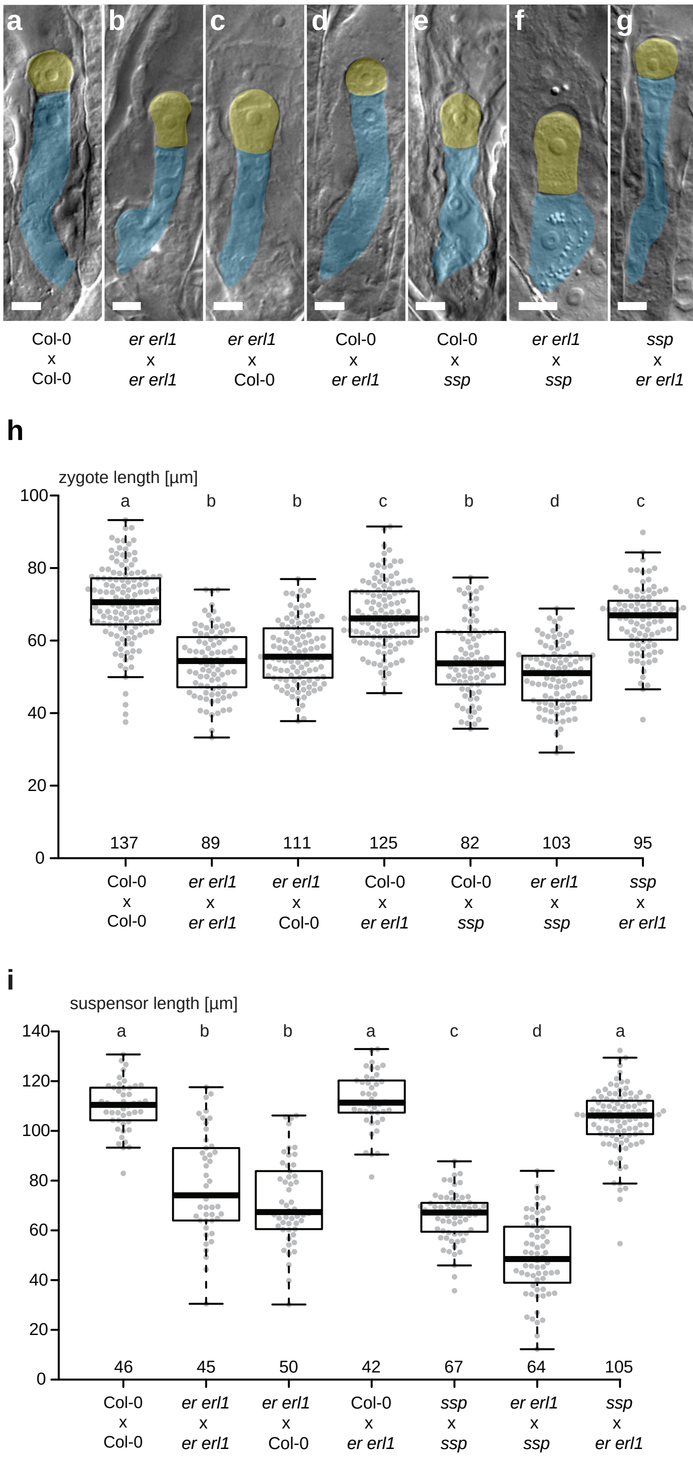

**Figure S4: Sporophytic maternal effect of *er erl1* double mutant**

**a-g)** DIC images of cleared ovules showing representative 1-cell F1 embryos. Apical cells are false colored in yellow, basal cells in blue. The genotype is given below the image as maternal x paternal cross. Scale bar = 10 µm.

**h)** Box plot diagram of zygote length in µm. The sample size is given above x-axis. Center lines show the medians; box limits indicate the 25th and 75th percentiles; whiskers extend 1.5 times the interquartile range from the 25th and 75th percentiles; small crosses represent sample means; data points are plotted as grey dots. Statistical differences determined by Mann-Whitney U-test ($p<0.05$) are indicated by different letters above graph.

**i)** Box plot diagram of suspensor length of transition stage embryos in µm. The sample size is given above x-axis. Center lines show the medians; box limits indicate the 25th and 75th percentiles; whiskers extend 1.5 times the interquartile range from the 25th and 75th percentiles; small crosses represent sample means; data points are plotted as grey dots. Statistical differences determined by Mann-Whitney U-test ($p<0.05$) are indicated by different letters above graph.

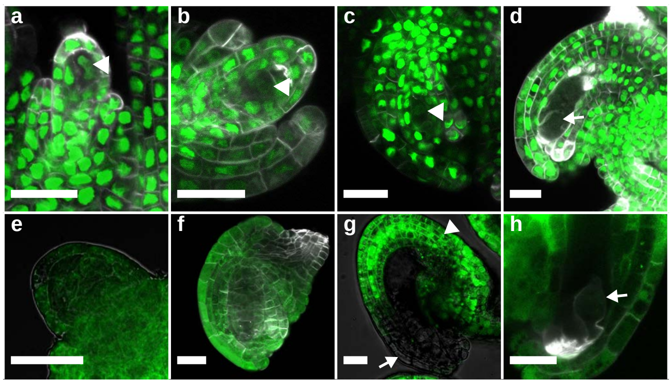

**Figure S5: *ERECTA* expression in developing ovules**

**a-d)** Confocal microscopy of transcriptional reporter *pER::3xVenus-N7* in developing ovules. Nuclear-localized Venus YFP signal can be observed in the megaspore mother cell (arrowhead; panel **a**), the functional megaspore (arrowhead, panel **b**), the developing female gametophyte (arrowhead, panel **c**). In mature egg cells (arrow, panel **d**) the signal is too weak to be confidently identified.

**e-h)** Confocal microscopy of translational fusion reporter *pER::ER-YPet* in developing ovules. YPet YFP signal can be observed in the megaspore mother cell **e)**, the developing female gametophyte **f)**. Strong signal can also be observed in the integuments (arrowhead) of ovules with mature female gametophyte. In mature egg cells (arrow, panel **h**) the signal is too weak to be confidently identified.

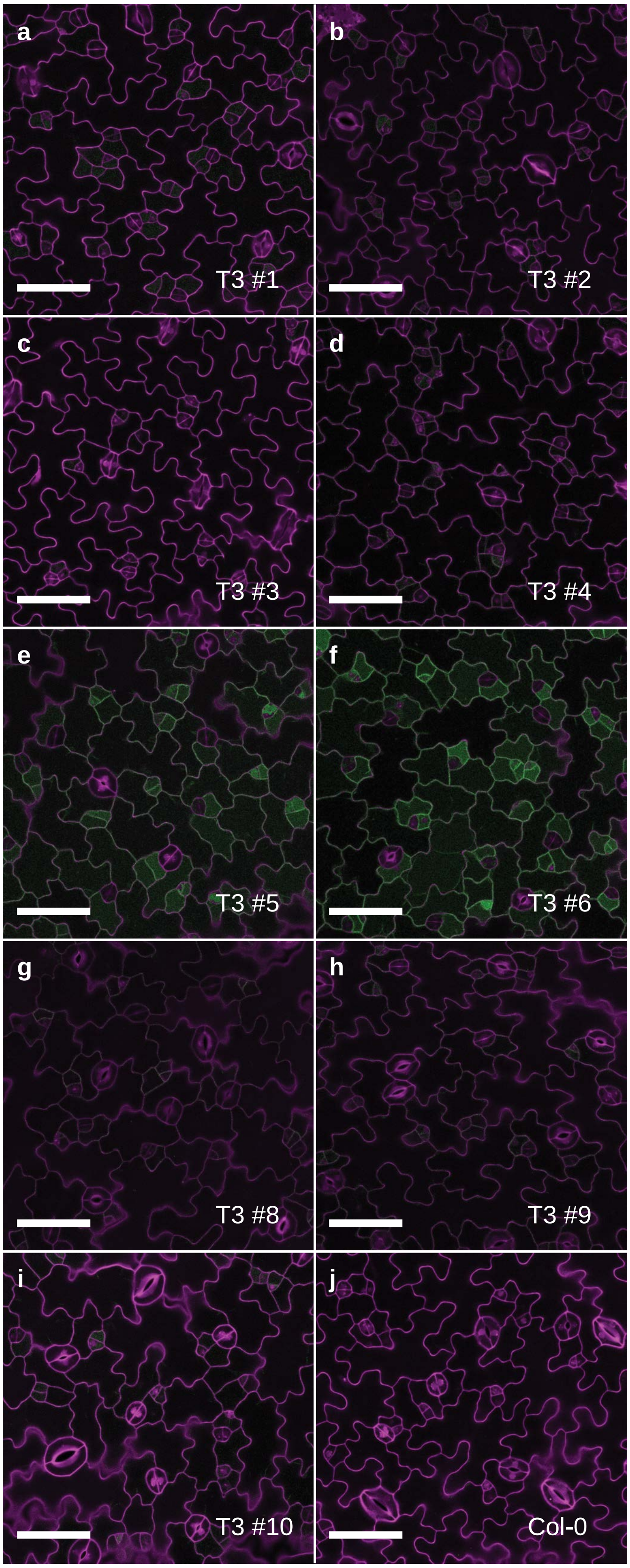

**Figure S6: Expression of *pER::ER-YPet* in *er erl1 erl2***
Genetic rescue by expression of *pER::ER-YPet* in *er erl1 erl2* mutants. Maximum projections of confocal image stacks of abaxial leaf surfaces of T3 seedlings 5 days after germination. YPet YFP fluorescence shown in green, propidium iodide staining shown in magenta. **a-j)** independent transgenic lines, **j)** wildtype. Scale bar = 50 µm.

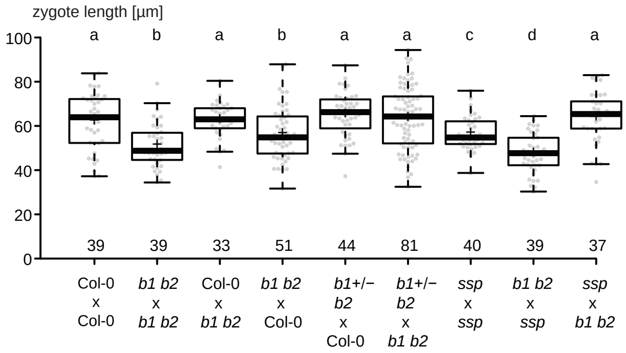

**Figure S7: Sporophytic maternal effect of *bsk1 bsk2* double mutants**

Box plot diagram of zygote length in µm of F1 embryos of crosses (maternal x paternal). Heterozygous mutations are indicated by +/-; *b1* = *bsk1*, *b2* = *bsk2*. The number of analyzed zygotes is given above x-axis. Center lines show the medians; box limits indicate the 25th and 75th percentiles; whiskers extend 1.5 times the interquartile range from the 25th and 75th percentiles; small crosses represent sample means; individual data points are plotted as grey dots. Statistical differences determined by Mann-Whitney U-test ($p<0.05$) are indicated by different letters above graph.

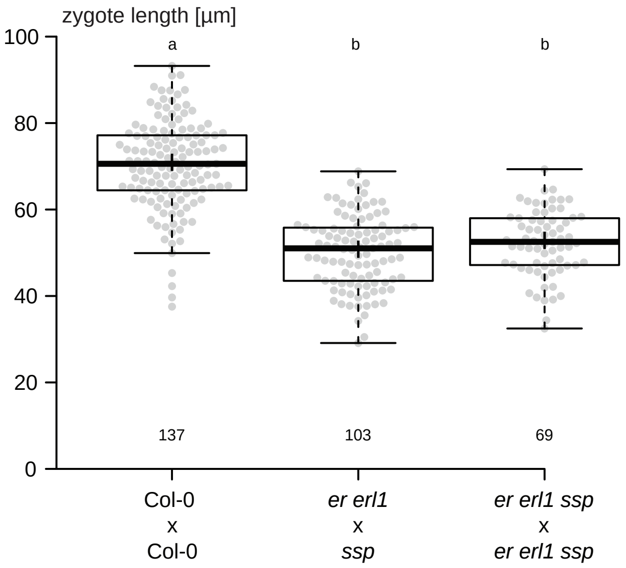

**Figure S8: Parental effects of *er erl1* and *ssp* mutations**
Box plot diagram of zygote length in µm of F1 embryos of crosses (maternal x paternal). The maternal effect of the *er erl1* double mutant and the paternal effect of the *ssp* mutant are additive in F1 embryos and are indistinguishable from the phenotype of self-pollinated *er erl1 ssp* triple mutants. The number of analyzed zygotes is given above x-axis. Center lines show the medians; box limits indicate the 25th and 75th percentiles; whiskers extend 1.5 times the interquartile range from the 25th and 75th percentiles; small crosses represent sample means; individual data points are plotted as grey dots. Statistically significant differences determined by Mann-Whitney U-test ($p<0.05$) are indicated by different letters above graph.

| | mRNA abundance | | | | |
|---|---|---|---|---|---|
| Gene name | EC | Z14h | Z24h | 1CE | 32CE |
| *ERECTA* | 70.87 | 25.50 | 8.08 | 5.19 | 15.89 |
| *ERL1* | 1.13 | 1.01 | 1.24 | 9.71 | 61.10 |
| *ERL2* | 1.32 | 1.89 | 3.40 | 33.76 | 69.86 |

**Supplementary Table S1: mRNA abundance of *ERECTA* family genes during early embryogenesis**
Expression level (FPKM value) of isolated egg cells (EC), zygotes 14 hours after pollination (Z14h), zygotes 24 hours after pollination (Z24h), 1-cell embryos (1CE), and globular stage embryos (32CE) derived from Zhao et al. (2019).

| Parental genotype | Number of seedlings triple mutant | total | Frequency [%] | Chi Square | P value |
|---|---|---|---|---|---|
| er+/- erl1 erl2 | 33 | 139 | 23.74 | 0.0583 | 0.809 |
| er erl1+/- erl2 | 38 | 150 | 25.33 | 0.0045 | 0.947 |

**Supplementary Table S2: Segregation rates of *er* and *erl1* in triple mutant combinations**
Segregation rate of triple homozygous offspring of *er +/- erl1 erl2* and *er erl1+/- erl2*. The frequency of triple homozygous seedlings does not deviate significantly from 25%, expected for normal Mendelian segregation (Chi Square test, $p<0.05$).

| Gene name | Gene ID | half life1 | half life2 | half life3 | Mean | SD | CI min | CI max |
|---|---|---|---|---|---|---|---|---|
| ER | AT2G26330 | 2.32 | 1.60 | 1.42 | 1.78 | 0.39 | 1.56 | 2.00 |
| ERL1 | AT5G62230 | 3.97 | 2.46 | 0.68 | 2.37 | 1.35 | 1.60 | 3.15 |
| ERL2 | AT5G07180 | 4.18 | 1.45 | 1.88 | 2.50 | 1.20 | 1.81 | 3.19 |
| BSK1 | AT4G35230 | 1.46 | 1.24 | 0.84 | 1.18 | 0.26 | 1.03 | 1.33 |
| BSK2 | AT5G46570 | 2.77 | 1.40 | 2.58 | 2.25 | 0.61 | 1.90 | 2.60 |
| YDA | AT1G63700 | 1.33 | 0.66 | 0.72 | 0.91 | 0.30 | 0.73 | 1.08 |
| MPK6 | AT2G43790 | 2.87 | 1.44 | 1.32 | 1.88 | 0.70 | 1.48 | 2.28 |
| FLS2 | AT5G46330 | 1.18 | 0.53 | 0.52 | 0.74 | 0.31 | 0.57 | 0.92 |
| BRI1 | AT4G39400 | 1.06 | 0.65 | 0.67 | 0.79 | 0.19 | 0.68 | 0.90 |
| HAE | AT4G28490 | 1.73 | 0.50 | 0.84 | 1.02 | 0.52 | 0.73 | 1.32 |
| ACT2 | AT3G18780 | 2.68 | 1.82 | 1.74 | 2.08 | 0.42 | 1.84 | 2.33 |

**Supplementary Table S3: mRNA turn-over of selected candidate transcripts in Arabidopsis seedlings**
mRNA half life in hours; mean values of three replicates, standrad deviation (SD), and minimal and maximal values of confidence interval (CI) are given

**Supplementary Table S4: Primer sequences**

Genotyping primers

| mutant | LP primer | RP primer |
|---|---|---|
| er | TTCGAAATCGAAAACGGTATG | TGTGTGTGAGAAATGGCTCTG |
| erl1 | TTTCCAATCATGATGTTGCAG | CAAACAATTGCTCCAGCTTTC |
| erl2 | TATCTCCATGGCAACAAGCTC | AATGACACATCGCTGAGAAGG |
| yda-mb1 | GGGTTGTTGATTAGTAAACCATAT | TAGTAGGAGACCCAGTAGTT |
| ssp-2 | TTAGAGACCACACGAGAAGGC | TAACATGGCTTGGTCTGATCC |
| bsk1-2 | ATGAGGTTGCGAGTAGGAT | CAAAAGCATGACCAATGAGAAG |
| bsk2-2 | GGAAGCGACTGGTGTTGGGAAG | TGGTCGATCCTTTGCCTCTGA |

LBb1.3 for SALK lines:
ATTTTGCCGATTTCGGAAC
GABI-LB for GK lines:
ATATTGACCATCATACTCATTGC

Primers for plasmid construction

pERECTA-IF-1F
TTCGAGCTCGGTCCCGGGCAAGATCCTGGATTTGTAACTG
pERECTA-IF-1R
TCACCATGGTGGATCCttctcacacacagtcttaaaacgac
pERECTA-IF-2F
ataaaataatgtcgacCAAGATCCTGGATTTGTAACTG
ERECTA-kinase-IF-R
TTTAGACACCATCCCCTCACTGTTCTGAGAAATAACTTGTCCAAAC
Nanobody-IF-F:
TAGTAAAAAAGGATCCATGATGAAAATGGAGACTGAC
Nanobody-IF-R:
TGGTTACCTTTTTAGGGCCCTTAGCTGGAGACGGTGACCTGG
pEC1.1-IF-F:
ataaaataatgtcgacCGCCTTATGATTTCTTCGGTTTC
pEC1.1-IF-R:
ATCCTTTTTTACTAGTCCTTCTCAACAGATTGATAAGG

**Material and Methods**

Plant lines and growth conditions

All *Arabidopsis thaliana* plants used in this study were grown under long-day conditions as described before [1]. The *ssp-2* single[2], *bsk1 bsk2* double mutant[3], *er* (SALK_066455), *erl1* (GK_109G04), *erl2* (GK_486E03)[3] have been described previously. The *yda-10* (SALKseq_078777) allele in Col-0 background carries a T-DNA insertion in the second exon. As the truncated gene product lacks essential parts of the YDA coding region[4] and the mutant phenotype is similar to *yda-1* [4], the *yda-10* presumably represents a null allele. Insertion lines were kindly provided by the Nottingham Arabidopsis Stock Center [5]. Multiple mutant combination were obtained by crossing. Crosses were performed by manual dissection of anthers before anthesis, followed by manual pollination approx. 24h later.

Genotyping

For genotyping of mutant plants, PCR was performed with ThermoFisher DreamTaq DNA polymerase. Gene-specific primers (LP and RP; Table S4) were used for the wild-type allele. Left border primers in combination with a gene-specific primer (RP) were used to detect the insertion allele.

Plasmid construction

Plasmid construction was performed by in-fusion cloning (TaKaRa) according to the manufacturer's recommendation. Oligonucleotide sequences for molecular cloning are summarized in Table S4. For plant transformation, transgenes were constructed in pCambia 1300 (GenBank: AF234296) and pCambia 3300 as well as pBay-bar and pBay-hyg binary vectors. pBay-bar and pBay-hyg are modified versions of the pCambia binary vectors, where the T-DNA has been replaced by a synthetic DNA fragment harboring unique restriction enzyme sites as well as plant codon-optimized selectable marker genes (conferring phosphinothricin and hygromycin, respectively) under control of the Arabidopsis *RPL10A* promoter (-960 to +18 including the first intron of At1g14320). A 1934bp region immediately upstream of the start codon of *ER* was transcriptionally fused with the coding region of three copies of Venus YFP with a C-terminal N7 nuclear localization sequence in pBay-bar to generate *pER::3xVenus-N7.* pBay-bar *pER::ER-YPet* contain a fragment of the *ER* locus including a 1934bp region upstream of the start codon and the entire protein-coding region as well as all introns. The *ER* genomic sequence was fused in-frame at the 3' end with the coding region of *YPet YFP*. To generate *pER::NSlmb-vhhGFP4,* the ER-YPet coding region in pBay-bar *pER::ER-YPet* was replaced by the *NSlmb-vhhGFP4* coding region [6] and the resulting chimeric gene transferred to pBay-hyg. For *pEC1::NSlmb-vhhGFP4,* a 465bp region of *EC1.1* was fused with *NSlmb-vhhGFP4* in pBay-hyg. *p35S::SSP-YFP*, *pSSP::SSP-YFP, and pYDA::yda-CA* have been described previously [2,4]

Plant transformation

All transgenic plants were transformed by floral dip using *Agrobacterium tumefaciens* GV3101. *pER::3xVenus-N7* was transformed into Col-0, *pER::ER-YPet* and *p35S::SSP-YFP* into *er erl1+/- erl2* plants. *pYDA::yda-CA* and *pSSP::SSP-YFP* were transformed into *er erl2* and *er erl1*, respectively. Transgenic seeds were screened on ½ MS plates containing 50gm/L phosphinothricin. T3 plants were used for phenotypic analyses, except for *p35S::SSP-YFP,* where T1 plants were used because of sterility of the transgenic plants. Two independent transgenic lines of *pYDA::yda-CA* in *er erl2* (#16,

#19) were crossed with Col-0 and offspring homozygous for the *ER ERL2* wild-type alleles selected in the F2 generation. *pER::NSlmb-vhhGFP4* and *pEC1::NSlmb-vhhGFP4* were transformed into *pER::ER-YPet* in *er erl1 erl2*. T1 seedlings were screened on ½ MS plates containing 50 mg/L basta and 20 mg/L hygromycin.

Microscopy

For zygote length and suspensor length measurement, immature ovules were dissected by hand and incubated overnight in Hoyer's solution. Differential interference contrast (DIC) images were taken with a Zeiss Axio Imager.Z1 microscope equipped with AxioCam HRc camera and AxioVision LE software as described before [3]. Size measurements were performed using measurement tools of ImageJ software [7]. Zygote length was determined in fully elongated zygotes after the first zygotic cell division as a sum of apical and basal daughter cell size. Zygote polarity was determined as ratio of apical and basal cell sizes. Suspensor length during transition stage was measured from the micropylar end of the basal suspensor cell to the center of uppermost suspensor cell. Confocal microscopy was conducted with a Zeiss LSM 780 NLO microscope with ZEN software. SCRI Renaissance 2200 (SR2200) staining was performed according to published protocols [8,9] and confocal images were obtained with excitation at 405nm and detection wavelength at 415nm to 475nm. For YPet and Venus YFP, a 514nm laser wavelength was used for excitation, a wavelength between 526nm and 553nm was recorded. For propidium iodide (PI) staining, cotyledons of seedlings (5 days after germination) were dissected and incubated in 10mg/L PI solution in water for 30 minutes, followed by brief washing with water. Afterwards, cotyledons were transferred to microscopy slides. PI fluorescence was detected at 571nm to 656nm with an excitation wavelength of 561nm.

Phenotypic analysis of rosette leaves

For Images of rosette leaves, three-week old plants were photographed with Canon EOS 1000D camera. To increase the visual contrast, images of rosette leaves were isolated from background soil using Adobe Photoshop.

RNA stability

Arabidopsis RNA half-live (stability) was determined non-invasively in whole seedlings and results have been published recently [10].

Data visualization and statistical analysis

Box plot diagrams were made with BoxPlotR [11]. Mann-Whitney U-test was used for statistical analysis of phenotypic data. The segregation analysis was statistically analyzed with Chi Square test.

1 Babu, Y., Musielak, T., Henschen, A. & Bayer, M. Suspensor Length Determines Developmental Progression of the Embryo in Arabidopsis. *Plant Physiology* **162**, 1448-1458, doi:DOI 10.1104/pp.113.217166 (2013).

2 Bayer, M. *et al.* Paternal control of embryonic patterning in Arabidopsis thaliana. *Science* **323**, 1485-1488, doi:323/5920/1485 [pii] 10.1126/science.1167784 (2009).

3 Neu, A. *et al.* Constitutive signaling activity of a receptor-associated protein links fertilization with embryonic patterning in Arabidopsis thaliana. *Proc Natl Acad Sci U S A* **116**, 5795-5804, doi:10.1073/pnas.1815866116 (2019).

4 Lukowitz, W., Roeder, A., Parmenter, D. & Somerville, C. A MAPKK kinase gene regulates extra-embryonic cell fate in Arabidopsis. *Cell* **116**, 109-119, doi:S0092867403010675 [pii] (2004).
5 Scholl, R. L., May, S. T. & Ware, D. H. Seed and molecular resources for Arabidopsis. *Plant Physiol* **124**, 1477-1480, doi:10.1104/pp.124.4.1477 (2000).
6 Ma, Y. *et al.* WUSCHEL acts as an auxin response rheostat to maintain apical stem cells in Arabidopsis. *Nat Commun* **10**, 5093, doi:10.1038/s41467-019-13074-9 (2019).
7 Schneider, C. A., Rasband, W. S. & Eliceiri, K. W. NIH Image to ImageJ: 25 years of image analysis. *Nat Methods* **9**, 671-675 (2012).
8 Musielak, T. J., Schenkel, L., Kolb, M., Henschen, A. & Bayer, M. A simple and versatile cell wall staining protocol to study plant reproduction. *Plant Reprod* **28**, 161-169, doi:10.1007/s00497-015-0267-1 (2015).
9 Musielak, T. J., Bürgel, P., Kolb, M. & Bayer, M. Use of SCRI Renaissance 2200 (SR2200) as a Versatile Dye for Imaging of Developing Embryos, Whole Ovules, Pollen Tubes and Roots. . *Bio-protocol* **6**, e1935, doi:10.21769/BioProtoc.1935 (2016).
10 Szabo, E. X. *et al.* Metabolic Labeling of RNAs Uncovers Hidden Features and Dynamics of the Arabidopsis Transcriptome. *Plant Cell* **32**, 871-887, doi:10.1105/tpc.19.00214 (2020).
11 Spitzer, M., Wildenhain, J., Rappsilber, J. & Tyers, M. BoxPlotR: a web tool for generation of box plots. *Nat Methods* **11**, 121-122, doi:10.1038/nmeth.2811 (2014).
